# Does age protect against loss of tonotopy after acute deafness in adulthood?

**DOI:** 10.1101/2024.05.03.592347

**Authors:** Nicole Rosskothen-Kuhl, Sarah Green, Till F. Jakob

## Abstract

The mammalian auditory system develops a topographical organization of sound frequencies along its pathways, also called tonotopy, as a result of early auditory input. In contrast, sensory deprivation during early development results in no or only rudimentary tonotopic organization. This study addresses two questions: 1) How robust is the tonotopy when hearing fails in adulthood? 2) What role does age play at time of deafness? To address these questions, we deafened young and old adult rats with normal hearing. One month after deafness, both groups were unilaterally supplied with cochlear implants and electrically stimulated for two hours. The central auditory neurons, which were activated as a result of the local electrical intracochlear stimulation, were visualized using Fos staining. While the auditory system of young rats lost the tonotopic organization throughout the brainstem, the auditory system of the older rats mainly sustained its tonotopy. It can be proposed that neuronal plasticity prevails in the central auditory system of young adult rats, while neuronal stability prevails in the brains of aging rats. Consequently, age may be an important factor in protecting a hearing-experienced adult auditory system from a rapid loss of tonotopy when suffering from acute hearing loss. Furthermore, the study provides compelling evidence that acute deafness in young adult patients should be diagnosed as early as possible to prevent maladaptation of the central auditory system and thus achieve the optimal hearing outcome with a hearing prosthesis.

## 1. Introduction

Recently around 5% of the world’s population (∼430 million people) suffer from disabling hearing loss (WHO, 2024) making it the most common sensory impairment of our age. The majority of these are adults (∼396 million). The prevalence of hearing loss increases with age, reaching approximately 25% for the population over 60 years of age, with an upward trend (Roth et al., 2011; Gablenz et al., 2017; WHO, 2024). Age-related hearing loss is mostly caused by loss of hair cells and is often associated with changes in the auditory nerve and the central auditory system, among other factors (Willott, 1984; Palombi and Caspary, 1996; McFadden et al., 1997; Ingham et al., 1998; Syka, 2002; Ouda et al., 2015). Currently, this form of hearing loss is not reversible. Nevertheless, following the diagnosis of hearing loss patients can be (re)integrated into an acoustic environment through the use of hearing aids or neuroprostheses such as the cochlear implant (CI). For people with severe to profound sensorineural hearing loss, CIs can be highly beneficial, allowing for near-normal spoken language acquisition. For prelingually deaf patients best performances can be observed when CI implantation takes place during early development, specifically within the first three years of life (Kral and Sharma, 2012). In contrast, postlingually deaf CI patients who had normal hearing in early development can still achieve good hearing performances in adulthood, even after more than 10 to 30 years of deafness (Moon et al., 2014; Hiel et al., 2016; Medina et al., 2017; Kim et al., 2018) although they show better speech understanding with a duration of deafness less than 10 years (Kim et al., 2018). While Kim et al. (2018) observe better word recognition scores for the CI patients with a younger age at implantation, Garcia-Iza et al. (2018) suggest that the age at onset of postlingual deafness does not affect the CI outcome for patients with similar duration of deafness. Moon et al. (2014) have identified a positive effect on the performance of CI patients when deafness occurs after the age of 13 and according to Hiel et al. (2016), “age should not be a limiting factor for cochlear implantation decision” of postlingually deafened patients. To the best of our knowledge, it is still unclear why older CI patients with reduced or altered plasticity (Burke and Barnes, 2006; Apple et al., 2017; Foscarin et al., 2017) can achieve similar or even better hearing performance than younger CI patients with more plastic brains. For example, adult-deafened CI patients demonstrate significantly enhanced performances in sound source localization using interaural time differences, in comparison to prelingually deafened patients fitted with a CI at an early age, who often exhibit complete insensitivity to this cue (Litovsky et al., 2012; Ehlers et al., 2017).

The normal hearing central auditory system, from the cochlear nucleus to the auditory cortex, is organized tonotopically. This means that groups of neurons have the greatest sensitivity for different frequencies and are arranged spatially according to frequency. Auditory input during early development is required to shape auditory neurons into topographical maps. In the case of normal hearing, an organization of the auditory system can be observed already early in life (Kandler et al., 2009). In rats, an adult-like tonotopic arrangement matures before the third week of life (Friauf, 1992). However, previous studies have shown that tonotopy is altered in the absence of sensory input. In various model organisms, early bilateral hearing loss results in the reduction or even complete loss of tonotopic order in subcortical and cortical regions (Raggio and Schreiner, 1999; Leake et al., 2008; Fallon et al., 2009; Rosskothen-Kuhl and Illing, 2012; Jakob et al., 2015; Rauch et al., 2016; Jakob et al., 2019). In rats, we have demonstrated that unilateral or bilateral neonatal deafness results in the absence of tonotopic organization along the deprived auditory pathway (Rosskothen-Kuhl and Illing, 2012; Jakob et al., 2015; Rauch et al., 2016; Rosskothen-Kuhl et al., 2018; Jakob et al., 2019).

Aim of this study was to investigate on the neuronal level why the deafened auditory system of elderly is able to benefit from CI supply despite a reduced level of plasticity. We focused on the effect of deafness on the mature auditory brainstem and addressed two key questions: firstly, how stable is the tonotopic organization of a hearing-experienced auditory system when hearing fails in adulthood? and secondly, what role does age play at the time of deafness? To address these questions, we conducted a study examining tonotopic organization of the auditory brainstem in adult-deafened rats of different ages. The auditory system of deafened rats was re-activated by electrical intracochlear stimulation (EIS), and the stimulation-induced neuronal activity pattern was visualized by staining the plasticity and activity marker Fos in brain sections (Ehret and Fischer, 1991; Friauf, 1992; Guzowski et al., 2001; Jakob and Illing, 2009; Rapanelli et al., 2010; Rosskothen-Kuhl and Illing, 2010, 2012; Rauch et al., 2016; Rosskothen-Kuhl et al., 2018). While previous studies have demonstrated that local EIS of hearing-experienced rats results in a tonotopic activation of neurons along the auditory pathway according to the stimulation position, our work in hearing-inexperienced, deaf rats has shown that a comparable intracochlear stimulation leads to a significantly increased number of activated neurons and a broader spread of excitation over almost the entire auditory brainstem (Rosskothen-Kuhl and Illing, 2010, 2012; Jakob et al., 2015; Rauch et al., 2016; Rosskothen-Kuhl et al., 2018; Jakob et al., 2019). Building on our previous work, we were able to show here that the activity pattern of a hearing-experienced, adult-deafened auditory system varies greatly depending on the duration of hearing prior to onset of deafness and at the age at which deafness occurred. A reduction in tonotopic organization in the auditory system was correlated with a younger age at onset of deafness.

## 2. Materials and Methods

### 2.1 Experimental groups

Forty-seven female Wistar rats were divided into four groups: 1) young or old adult, normal hearing (NH) rats (n=15), 2) neonatally-deafened (ND), young adult rats (n=9), 3) young adult-deafened (YAD) rats (n=12), and 4) old adult-deafened (OAD) rats (n=11). The left cochlea of each rat was electrically stimulated for two hours. Additionally, two YAD rats and two OAD rats underwent cochlear implantation on their left side for two hours without stimulation and served as adult-deafened (AD) “implanted” controls. Further, four adult-deafened rats served as “pure” controls (YAD: n=2; OAD: n=2). At the time of perfusion, all rats in the “young” adult group were approximately four months old, whereas all rats in the “old” adult group were around 13 months old. ND animals and portions of the NH animals have already been utilized in previous studies and served as reference groups (Rosskothen-Kuhl and Illing, 2010, 2012). All procedures involving a total 55 rats were approved by the Regierungspräsidium Freiburg (permission number G-10/83). We confirm that all of our methods were performed in accordance with the relevant guidelines and regulations and that our study is reported in accordance with the ARRIVE guidelines.

### 2.2 Induction of adult deafness

Corresponding to Liu et al. (2011) three or 12 months old rats were systemically deafened by a combination of the loop diuretic ethacrynic acid (EA, REOMAX, Bioindustrial L.I.M. S.P.A., Italy, 50mg/20ml) and the antibiotic kanamycin (KM, Sigma-Aldrich, Germany, K4000). Anesthesia was initiated under 5 % isoflurane (Forene 100 % [V/V], Abbott GmbH & Co. KG, Germany) in an inhalation chamber and maintained with ∼1.5 % isoflurane while the rats were kept warm on a heating pad. Before placing a tail vein catheter, the tail of the rats was heated in 38°C warm water for vasodilatation. After rinsing the lateral tail vein with ca. 0.1 ml sterile 0.9 % NaCl solution, freshly prepared EA solution (75 mg/kg) was slowly injected into the tail vein. Afterwards, the tail vein was rinsed again with ca. 0.1-0.2 ml 0.9 % NaCl solution. In a second step, KM solution (500 mg/kg) was injected intramuscularly. To ensure a sufficient supply of liquid and electrolytes, Ringer’s solution was injected subcutaneously followed by subcutaneous Carprofen (4 mg/kg, Carprieve, Norbrook Laboratories Ltd., Northern Ireland) injection for pain relief. The co-administration of EA and KM results in a rapid and permanent hearing loss induced by irreversible hair cell lesion (Liu et al., 2011).

### 2.3 Verification of hearing function or hearing loss

Normal hearing or hearing loss due to pharmacological treatment was verified by measuring auditory brainstem responses (ABRs) as described in (Jakob et al., 2015; Jakob et al., 2016; Rosskothen-Kuhl et al., 2018; Buck et al., 2021, 2021) In short, under ketamine (80 mg/kg, Ketanest S, Medistar Arzneimittelvertrieb GmbH, Germany) and xylazine (12 mg/kg, Rompun, Bayer, Germany) anesthesia each ear was stimulated separately through hollow ear bars with 0.5 ms clicks (0.1–3 kHz) with peak amplitudes up to 95 dB SL. ABRs were recorded by averaging scalp potentials measured with subcutaneous needle electrodes between mastoids and the vertex of the rat’s head over 300 click presentations. While normal hearing rats typically exhibited click ABR thresholds near 0 dB SL, deafened rats showed increased hearing thresholds ≥95 dB SL to broadband click stimuli as well as pure tones (Rosskothen-Kuhl et al., 2021; Buck et al., 2023). Figures 1 A and B show ABRs of an adult-deafened rat before and after deafening by EA and KM injection. Additionally, all kanamycin treated rats consistently failed to show a motor response to a handclap. The absence of this so-called Preyer’s reflex indicates a sustained increase of ABR threshold above 81 dB SPL (Jero et al., 2001).

**Fig. 1:**
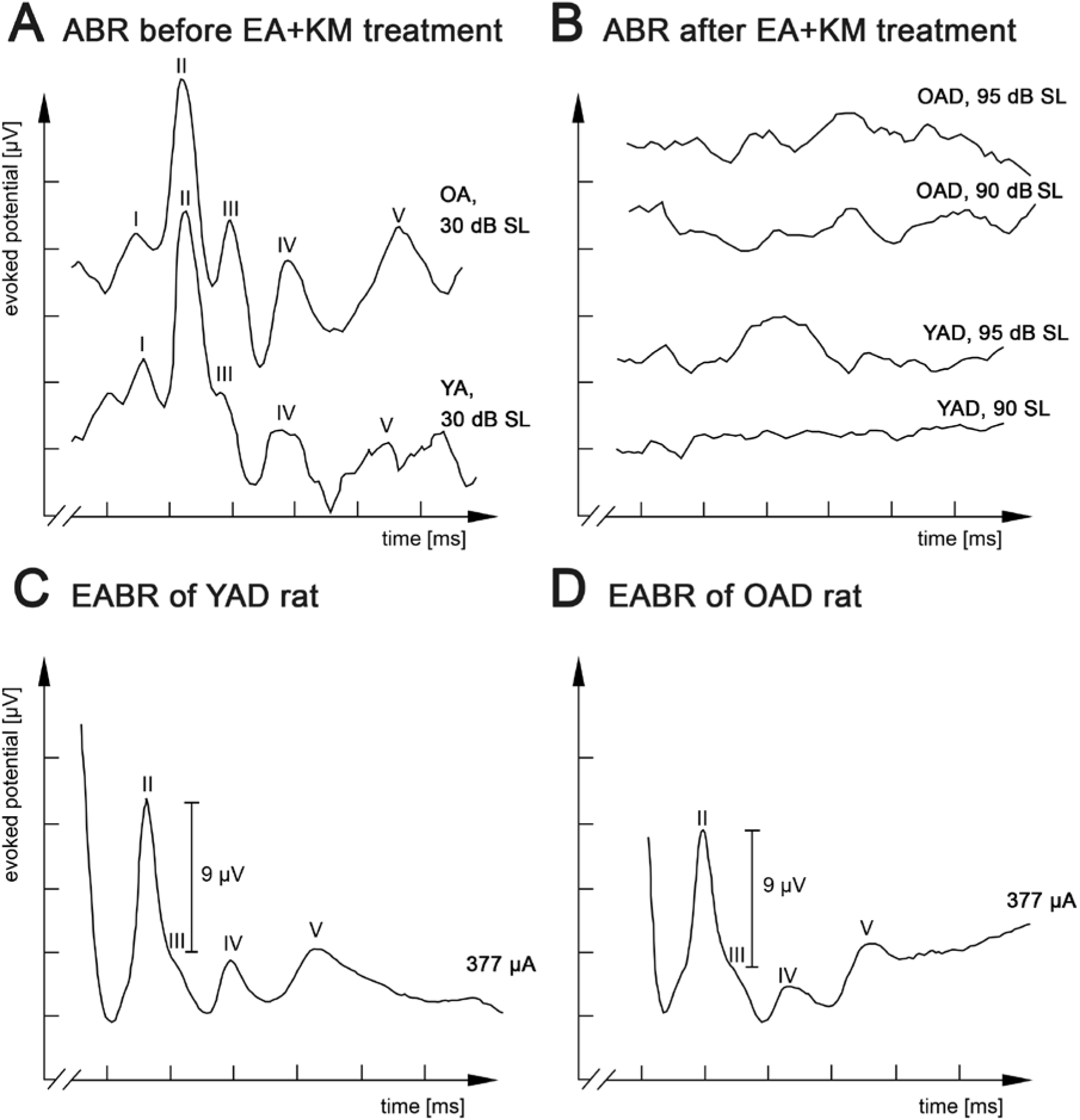
Kanamycin treatment in adult rats results in loss of auditory brainstem responses (ABR) and is restored by electrical intracochlear stimulation. A: Acoustic stimulation at 30 dB SL induced comparable ABRs in normally hearing old adult (OA) and young adult (YA) rats. B: ABR measurements of one young adult-deafened rat (YAD) and one old adult-deafened (OAD) rat after treatment with ethacrynic acid (EA) and kanamycin (KM). Electrical intracochlear stimulation at a current level of ∼377 µA induced comparable electrically evoked ABRs (EABRs) in YAD (C) and OAD (D) rats. I-V: recorded positive waves of acoustically or electrically evoked auditory brainstem potentials.

### 2.4 Electric intracochlear stimulation (EIS)

Two hours of EIS were applied under urethane anesthesia (i.p., 1.5 g/kg body weight; Fluka AG, Switzerland). In case of adult-deafened rats, unilateral EIS occurred on average 46 days after deafening. The stimulation set-up and cochlear implantation is described in detail in (Rosskothen-Kuhl and Illing, 2010, 2012, 2014; Jakob et al., 2015; Jakob et al., 2019). In short, two rings of an electrode array (CI24RE, Cochlear®, Australia) were inserted through a cochleostomy window over the middle turn of the rat cochlea corresponding to the 8-12 kHz region. The CI was connected to a Nucleus Implant Communicator kindly provided by Cochlear Germany GmbH and Co. KG. Electrically evoked ABRs (EABRs) were recorded to corroborate for the correct placement of stimulation electrodes and to determine an appropriate current level. Current levels for EIS were set to match an EABR amplitude of 9 µV±10%. This was achieved by a mean current level of ∼340 µA, corresponding to acoustic stimuli of around 85 dB above hearing threshold of NH rats. Corresponding to Rosskothen-Kuhl and Illing (2012), this stimulation intensity triggers tonotopic activation in the auditory system of NH rats.

### 2.5 Animal perfusion and preparation of brain sections

Following completion of the post-deafening survival time or stimulation period, rats were sacrificed by a lethal dose of sodium-thiopental (50 mg/ml per 200 g body weight of Trapanal 2.5 g, Nycomed, Germany) and perfused transcardially for 60 min. For perfusion, 4 °C cold fixative containing 4 % paraformaldehyde in 0.1 M phosphate buffer at pH 7.4 was used. Brains were removed from the skull and stored overnight in a phosphate buffer containing 20 % sucrose. Frontal brain sections of 30 µm thickness containing anteroventral cochlear nucleus (AVCN), lateral superior olive (LSO), and central inferior colliculus (CIC) were cut using a cryostat.

### 2.6 Immunohistochemistry

The immunostaining protocol that we used is described in detail in our previous studies (Illing et al., 2002; Rosskothen-Kuhl and Illing, 2010, 2012, 2014; Jakob et al., 2015; Jakob et al., 2019). Here, brain sections were exposed to a primary antibody raised in goat against Fos (SC-52-G, 1:2000, lot. no. L1406/ K1808/ F1109, Santa Cruz Biotechnology Inc., USA) (Rosskothen-Kuhl and Illing, 2010, 2012; Rosskothen-Kuhl et al., 2018). Visualization of primary antibody binding sites was based on the avidin–biotin technique (Cat. No. PK-6100, Vector Laboratories, USA), followed by staining with 0.05 % 3.3-diaminobenzidine tetrahydrochloride (Cat. No. 32750, Sigma, Germany), 0.3 % ammonium nickel (II) sulfate hexahydrate (Cat. No. A1827, Sigma, Germany) and 0.0015 % H_2_O_2_ in 50 mM Tris buffer.

### 2.7 Quantification of Fos-positive (+) nuclei

Photographs were taken from ipsilateral AVCN, ipsilateral LSO, and contralateral CIC of unilaterally stimulated rats using a 10x objective (for AVCN and LSO) or a 5x objective (for CIC) and a digital camera (Axiocam, Zeiss, Germany) at an 8-bit gray tone scale for quantitative evaluation of the staining results. Prior to the automated counting of Fos(+) nuclei, the color information of each image was removed by using Adobe Photoshop CS (Adobe Systems Inc., USA). Two regions of interest (ROI) were selected in AVCN, LSO, and CIC to determine differences in the spatial Fos expression pattern between the experimental groups (Fig. 6, red rectangles). While ROI 1 was placed in the mid-frequency range of NH rats, corresponding to intracochlear stimulation position, ROI 2 covered the low frequency region of NH rats (Ryan et al., 1988). Within the defined ROIs, gray value information was spread to the full 8-bit range (from 0 to 256). For automated counting of Fos(+) nuclei, photographs of three to five sections per rat through the anterior-posterior center of AVCN, LSO, and CIC were imported into the image analysis program iTEM (Olympus, Germany), where detection parameters were set as described in our previous studies (Rosskothen-Kuhl and Illing, 2010, 2012; Jakob et al., 2015; Jakob et al., 2019). The detection threshold for gray tone values was set to 145 for AVCN, 165 for LSO, and 200 for CIC. The ratio of ROI 2/ROI 1 was calculated for each brain section to identify differences in the stimulation-induced Fos expression pattern between the four different experimental groups (NH, ND, YAD, and OAD; Fig. 6). Ratios around 1 indicate a loss of tonotopy, while ratios closer to 0 indicate a more tonotopic organization of the corresponding auditory region.

### 2.8 Statistical Analysis

Statistical analysis was performed using Prism software (GraphPad Software, Inc., USA). We tested our data for normal distributions (using the Kolmogorov–Smirnov test) and equal variances (using the Brown–Forsythe and Bartlett tests). As our data did not consistently fulfill both criteria, we used the non-parametric Kruskal–Wallis and Mann–Whitney tests for statistical analysis. For both, significance level was set to p<0.05. In case of multiple comparisons, p-values were corrected using Dunn’s multiple comparisons test. All adjusted p-values as well as the number of values per group are reported in Tab. 1. In addition, we indicate the results of Kruskal–Wallis statistics (H) for each test. Significances were differentiated into (***) for p<0.001, (**) for p<0.01, and (*) for p<0.05 (Tab.1). Stereological corrections for counting particles in the sectioned material were not made as particle size was small compared to section thickness and comparisons are based on numerical relationships among objects of similar size rather than on absolute densities. The Fos analysis for AVCN, LSO, and CIC was based on a total of 47 rats.

**Tab. 1:**
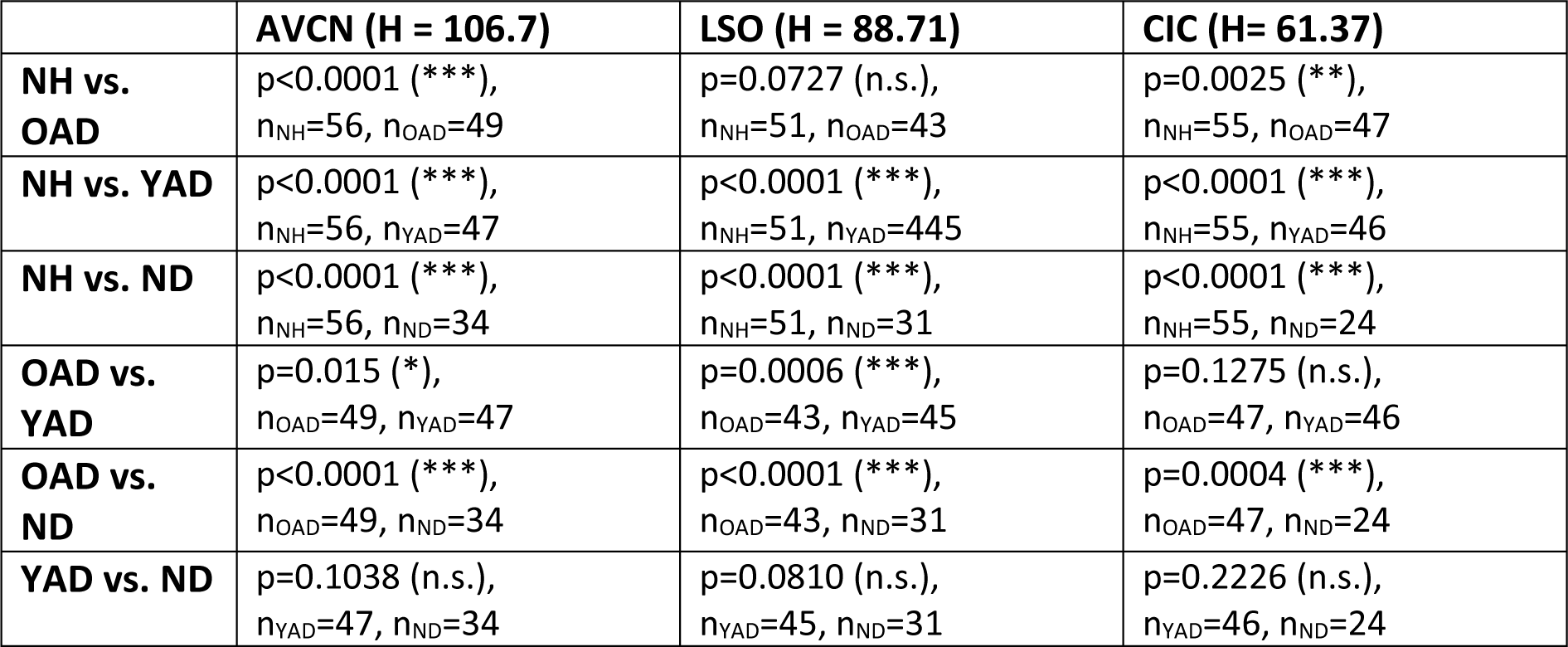
Statistical data of Fos quantification in three auditory brainstem regions of all four experimental groups: normally hearing (NH), neonatally-deafened (ND), young adult-deafened (YAD), old adult-deafened (OAD). H= Kruskal–Wallis statistics, n= number (n) of analyzed sections.

## 3. Results

### 3.1 Effect of kanamycin treatment in adult rats on auditory brainstem responses

To answer the research questions stated above, two cohorts of adult-deafened rats were studied: young adult-deafened (YAD) and old adult-deafened (OAD) rats. Before pharmacological treatment with kanamycin both cohorts showed normal hearing thresholds (young rats: x̅ = 14.3 dB SL; old rats: x̅ = 14.7 dB SL) and ABR responses with five distinguishable peaks (I–V, Fig. 1 A). Three days after EA+KM treatment, hearing thresholds increased by in mean 92 dB SL for young rats (n=16 rats) and 93 dB SL for old rats (n= 15 rats; Fig. 1 B). Despite hair cell loss the functionality of the auditory nerve has been preserved in both groups, which was verified by measuring EABRs with four well-differentiated peaks (II-V) under identical EIS. For both YAD and OAD rats, a peak II mean amplitude of 9 µV ± 10 % was achieved by applying a current of in mean ∼340 µA (Fig. 1C+D).

### 3.2 Deafening of young but not of old adult rats results in reduction of tonotopic organization

Deafness-induced changes in the tonotopic organization along the auditory brainstem were studied by staining brain sections containing AVCN (Fig. 2), LSO (Fig. 3), and CIC (Fig. 4) for the activity and plasticity marker Fos. Previous studies have demonstrated that Fos is a suitable marker for mapping changes in the activity pattern of the central auditory system (Rosskothen-Kuhl and Illing, 2010, 2012, 2014; Jakob et al., 2015; Jakob et al., 2016; Rosskothen-Kuhl et al., 2018; Jakob et al., 2019). However, its expression in auditory neurons requires at least 30 min of acoustic or electrical stimulation (Rosskothen-Kuhl and Illing, 2012).

**Fig. 2:**
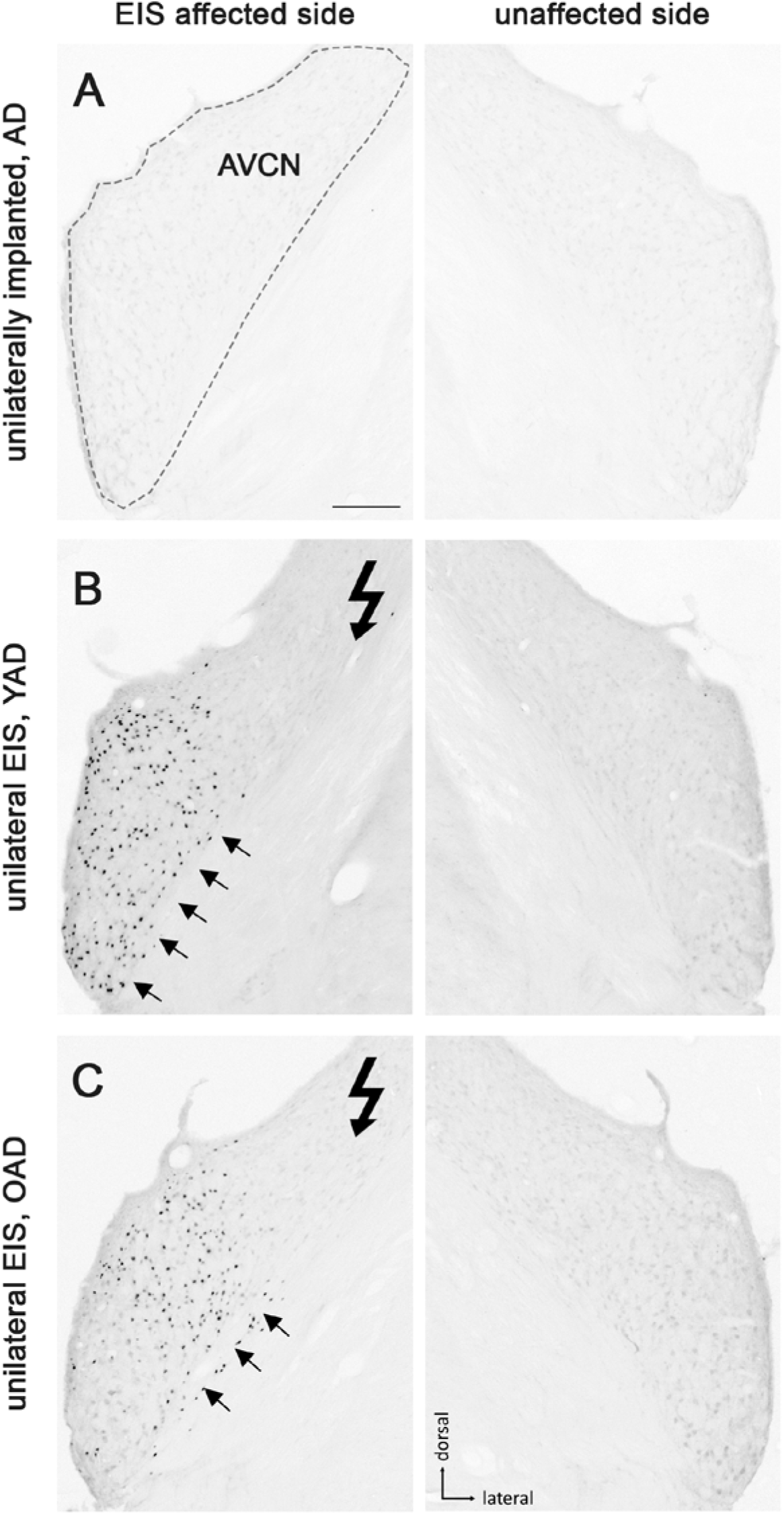
Patterns of Fos expression in the anteroventral cochlear nucleus (AVCN) of adult-deafened rats (AD, representative for the YAD and OAD cohort). A: No Fos expression was found in unstimulated but unilaterally implanted AD rats in both AVCNs. Dashed line shows the border of AVCN. B: Unilateral electrical intracochlear stimulation (EIS) of young adult-deafened (YAD) rats resulted in a large number of Fos expressing neurons (black dots) in the middle to ventral region of the ipsilateral AVCN (arrows), affected by EIS. C: Corresponding to the intracochlear electrode position, old adult-deafened (OAD) rats showed a Fos expression limited to the central region of the ipsilateral AVCN (arrows). Scale bar for A-C = 200 µm.

**Fig. 3:**
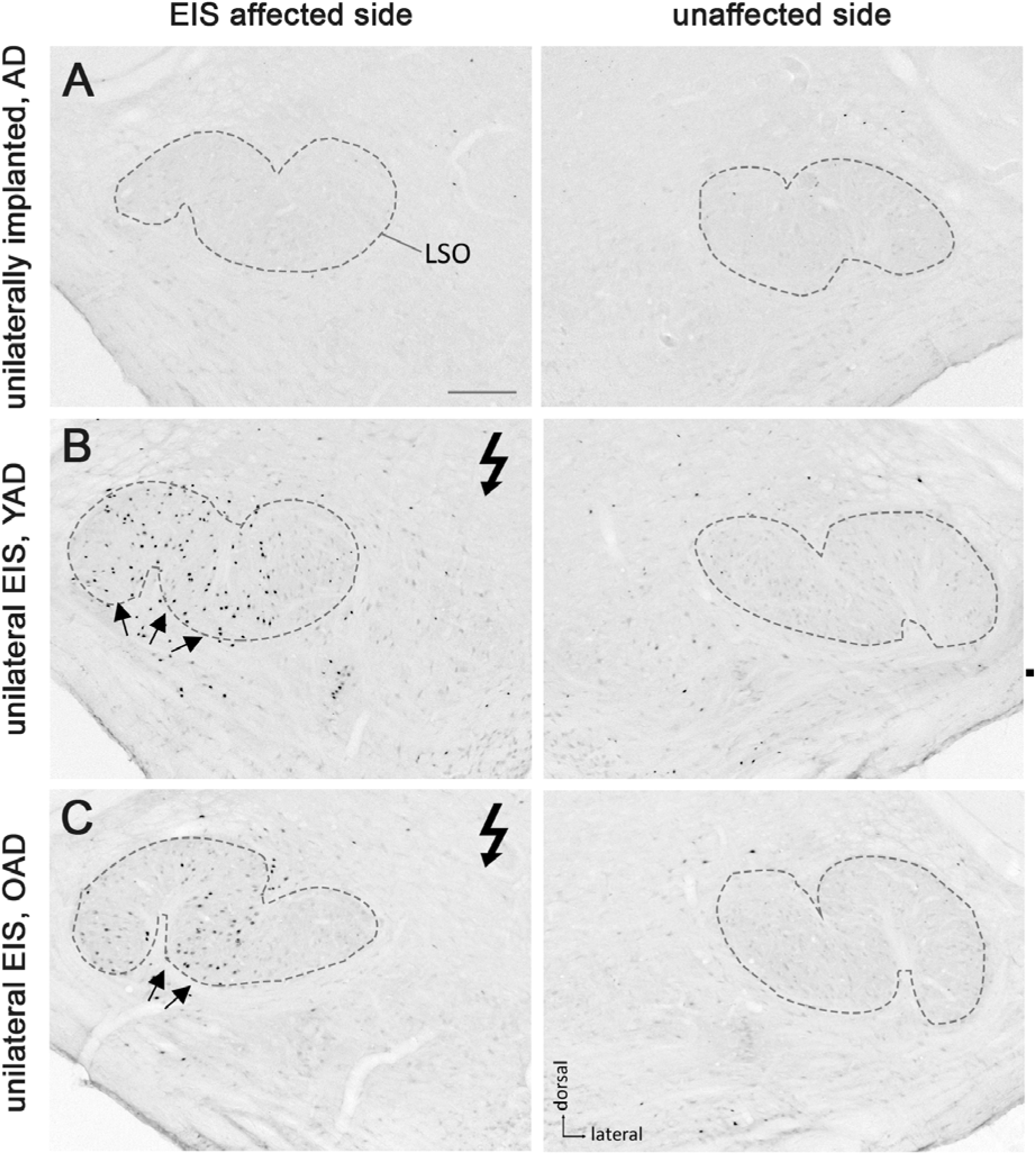
Patterns of Fos expression in the lateral superior olives (LSO) of adult-deafened rats (AD, representative for the YAD and OAD cohort). A: Marginal Fos expression was found in unstimulated but unilaterally implanted AD rats in both LSOs (dashed lines). B: Electrical intracochlear stimulation (EIS) of young adult-deafened (YAD) rats resulted in a large number of Fos expressing neurons (black dots) spread over nearly two-thirds of the EIS affected, ipsilateral LSO (arrows). C: Corresponding to the intracochlear electrode position, old adult-deafened (OAD) rats showed a Fos expression limited to the central region of the ipsilateral LSO (arrows). Scale bar for A-C = 200 µm.

**Fig. 4:**
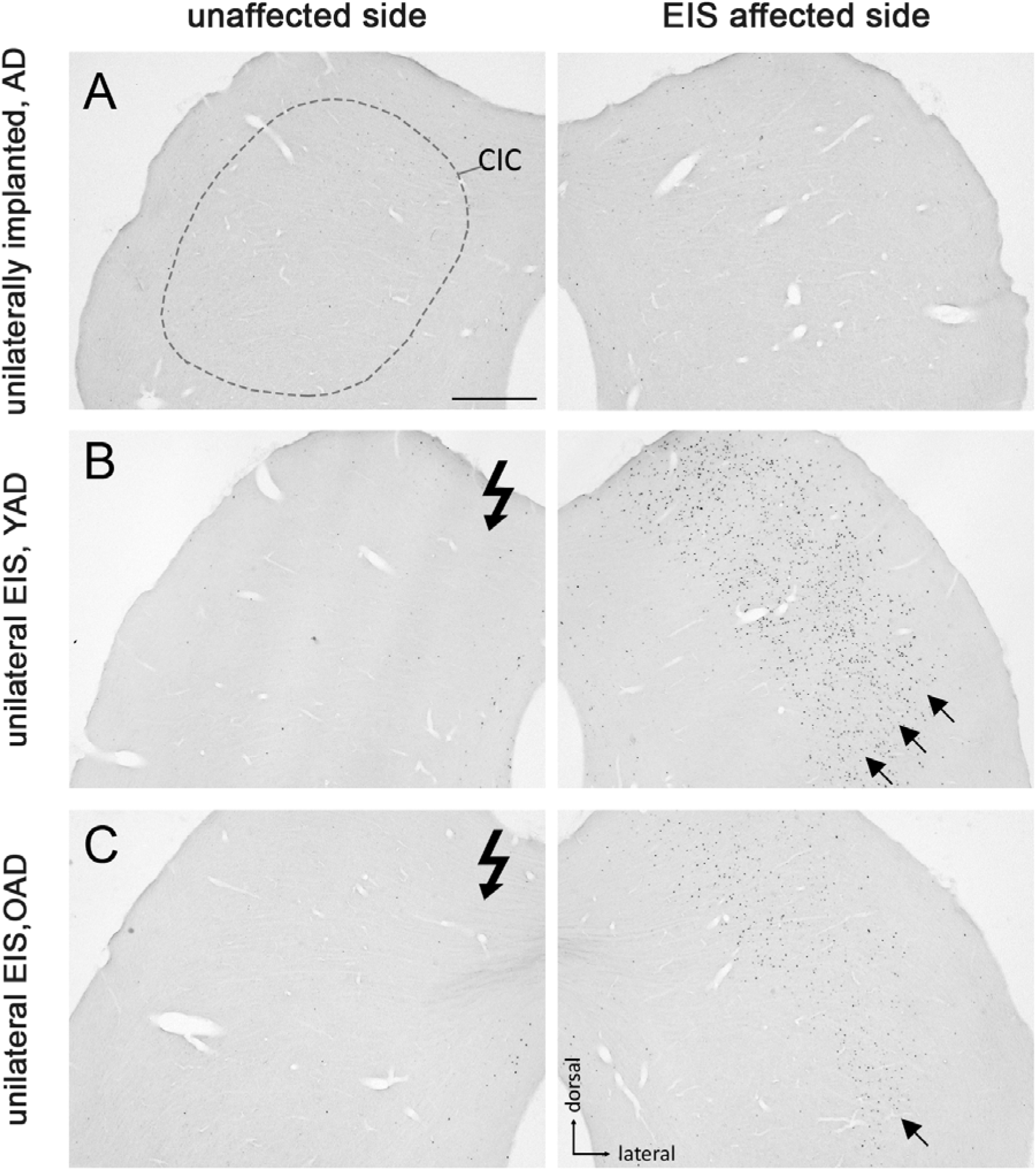
Fos expression pattern in the central inferior colliculus (CIC) of adult-deafened rats (AD, representative for the YAD and OAD cohort). A: Marginal Fos expression was found in unstimulated but unilaterally implanted AD rats in both CICs. Dashed line shows the border of CIC. B: Electrical intracochlear stimulation (EIS) of young adult-deafened (YAD) rats resulted in a large number of Fos expressing neurons (black dots) in the middle to lateral area of the contralateral CIC (arrows), affected by EIS. C: Corresponding to intracochlear electrode position, old adult-deafened (OAD) rats showed a Fos expression limited to the central region of the contralateral CIC (arrow). Scale bar for A-C = 500 µm.

One month after bilateral deafening of YAD and OAD rats, the mid-frequency region of their left cochlea was electrically stimulated for two hours. As a result, only the neurons of the stimulated auditory pathway (EIS affected side) expressed the Fos protein, although differences in the expression pattern were observed between the two cohorts. In YAD rats, a broadly distributed neuronal expression of Fos was observed over more than half of the ipsilateral AVCN (Fig. 2B arrows), the ipsilateral LSO (Fig. 3B arrows), and the contralateral CIC (Fig. 4B arrows) without clear boundaries indicating a tonotopic correspondence with the intracochlear stimulation position. Contralateral to stimulation (unaffected side), no Fos expression was observed in the AVCN, LSO, and CIC of this cohort (Figs. 2-4 B), which was consistent with the Fos expression pattern of implanted, non-stimulated controls (Figs. 2-4 A). In contrast, OAD rats showed a lower and more focused tonotopic Fos expression in the ipsilateral AVCN and LSO as well as in the contralateral CIC, corresponding to the position of intracochlear stimulation and indicated by Fos(+) neurons in the middle area of these auditory regions (Figs. 2-4 C arrows). Contralateral to the electrical stimulation (unaffected side), OAD rats also showed no Fos expression (Figs. 2-4 C), corresponding to controls.

### 3.3 The age at onset of deafness affects the tonotopic organization of the auditory system

Figure 5 shows the neuronal Fos expression in the auditory brainstem (AVCN, LSO, and CIC) of NH, OAD, YAD, and ND rats after two hours of EIS. In previous studies we have demonstrated that the expression pattern of Fos and thus the stimulation-induced activity in the central auditory brainstem massively changes when hearing fails in early development (Rosskothen-Kuhl and Illing, 2010, 2012; Jakob et al., 2015; Rauch et al., 2016; Rosskothen-Kuhl et al., 2018; Jakob et al., 2019). While local intracochlear stimulation of young and old NH rats induces a tonotopic Fos expression in auditory brainstem nuclei (Ryan et al., 1988; Rosskothen et al., 2008; Jakob and Illing, 2009; Rosskothen-Kuhl and Illing, 2010; Rauch et al., 2016) (Fig. 5 A-C arrow(s)), local EIS of ND rats results in a spread of neuronal activity beyond the “normal” frequency borders, which is shown by a massive increase and dispersion of Fos-positive nuclei (Rosskothen-Kuhl and Illing, 2012; Jakob et al., 2015; Rauch et al., 2016; Rosskothen-Kuhl et al., 2018; Jakob et al., 2019) (Fig. 5 J-L arrows). Neonatally-deafened brains thus show a lack of tonotopic organization along the auditory pathway, which is most likely a result of missing sensory input during early development. When comparing the stimulation-induced Fos expression patterns of NH rats (Fig. 5 A-C) and ND rats (Fig. 5 J-L) with the expression patterns observed in young and old adult-deafened animals (Figure 5 D-I), it becomes clear that: first, the neuronal activity pattern of YAD rats (Fig. 5 G-I) is very similar to the pattern of ND rats, and second, OAD rats (Fig. 5 D-F) show a more tonotopic Fos expression which is similar to the expression pattern of NH animals. We conclude that in contrast to a young adult auditory system an older auditory system is able to maintain its tonotopic organization even after one month of hearing deprivation.

**Fig. 5:**
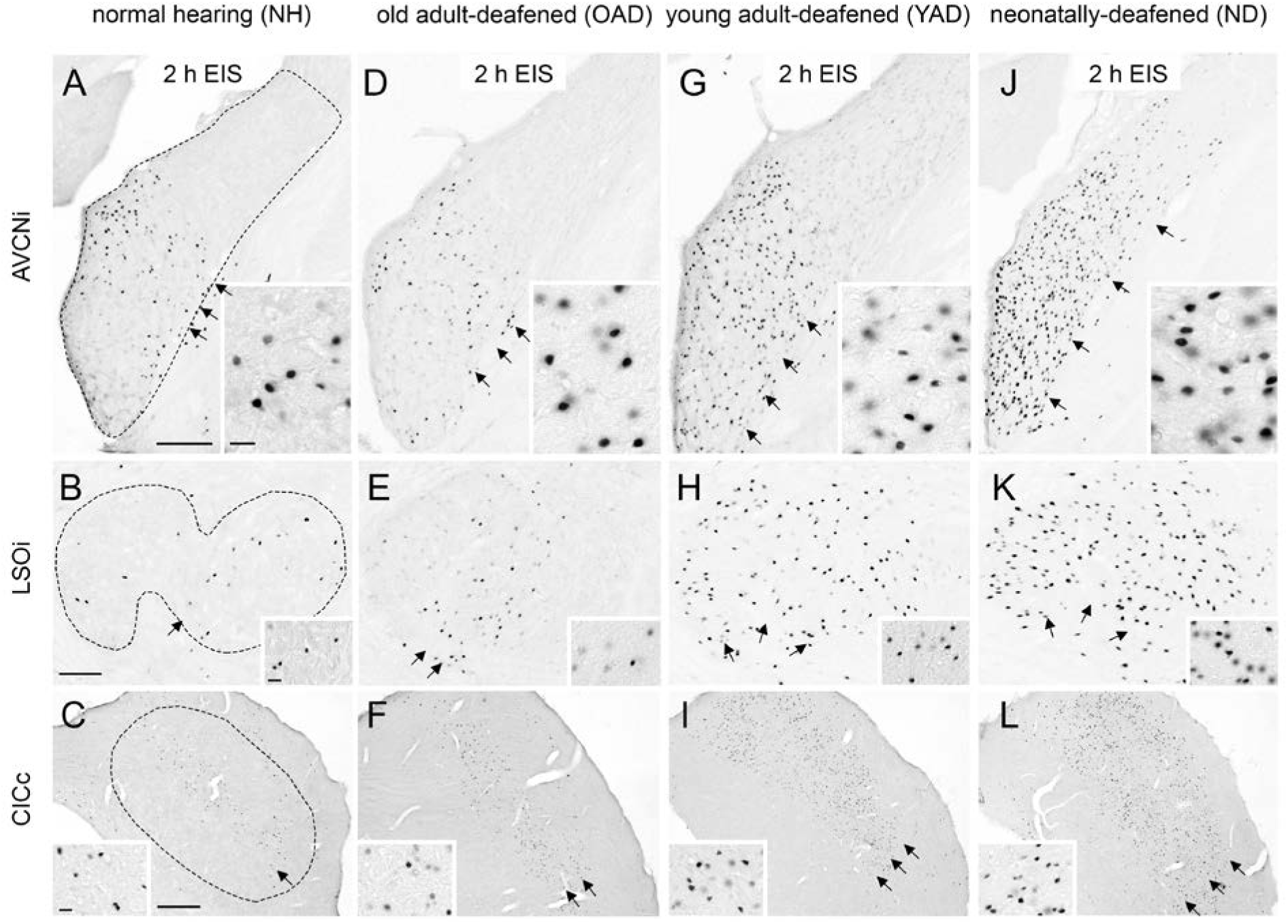
Stimulation-induced Fos expression patterns in neurons of the auditory brainstem of four experimental groups, normal hearing (NH), old adult-deafened (OAD), young adult-deafened (YAD), and neonatally-deafened (ND) rats. A-C: In young and old NH rats stimulation-induced Fos expression is found in a tonotopic order (arrows) in AVCNi (A), LSOi (B), and CICc (C). This corresponds to the intracochlear stimulation position in the mid-frequency range. D-F: Similar to NH rats, OAD rats showed a Fos expression with a tonotopic distribution (arrows) in the auditory brainstem. G-I: In contrast, YAD rats showed an increased and spread Fos expression pattern (arrows) indicating a degraded tonotopic organization. J-L: In the auditory brainstem of ND rats, Fos expression pattern (arrows) was similar to the pattern of YAD rats, recognizable by an increased number of widely spread Fos-positive neurons in the auditory brainstem regions. Insets show Fos-positive nuclei at 100x magnification. Scale bar = 20 µm. AVCNi: anteroventral cochlear nucleus ipsilateral to electrical stimulation; CICc: central inferior colliculus contralateral to electrical stimulation; LSOi: lateral superior olive ipsilateral to electrical stimulation. Scale bars in A and B = 200 µm, and in C = 500 µm.

### 3.4 The shorter the hearing experience before deafness, the less the development or preservation of tonotopic organization in the auditory brainstem

To identify a correlation between tonotopic organization of the auditory system and hearing experience prior to deafness, the topography of stimulation-induced Fos expression in AVCN (Fig. 6 A, dashed line), LSO (Fig. 6 B, dashed line), and CIC (Fig. 6 C, dashed line) was determined for all four experimental groups: (1) young and old NH rats, (2) OAD rats, (3) YAD rats, and (4) ND rats. Defining two regions of interest (ROIs), with ROI 1 lying in the core region of mid-frequency sensitivity of NH rats (corresponding to our intracochlear stimulation position) and ROI 2 positioned in the region of lower frequency sensitivity of NH rats (Fig.6, red rectangles), the change of the local distribution of Fos(+) neurons was quantified by calculating the ratio of ROI 2/ROI 1. As a result, all three brainstem regions showed a significant increase of the ratio (ROI 2/ROI 1) with decrease of the hearing experience (Fig. 6; for p-values of statistical analysis see Tab.1). Comparing all four experimental groups (Fig. 6), it becomes clear that NH rats (young and old) showed the lowest ratio (median for AVCN=0.075, LSO=0.14, and CIC=0.4), indicating the highest degree of tonotopic order along their auditory brainstem followed by somewhat higher ratios for OAD rats (median for AVCN=0.34, LSO=0.4, and CIC=0.7). For all three auditory regions, the ratios of YAD rats were in median between 0.67 and 0.79 and thus higher than for the NH and OAD cohorts, which points out a massively reduced tonotopic order. The highest ROI 2/ROI 1 ratio with values close to 1 (Fig. 6, dashed line in statistical panels) were detected for the ND cohort (median for AVCN=0.82, LSO=1, and CIC=0.87), meaning that the distribution of Fos(+) neurons in both ROIs was almost equal and thus indicating a loss of tonotopy. In detail, for AVCN, we found significant differences between all groups, except for YAD and ND rats (Fig. 5 A; Tab. 1). In the LSO, we detected significant differences between groups, excluding NH vs. OAD rats and YAD vs. ND rats (Fig. 5 B; Tab. 1). For CIC, all groups showed significant differences between each other except YAD vs. OAD or YAD vs. ND rats (Fig. 5 C; Tab. 1). The detailed statistical results, including p-values, Kruskal–Wallis statistics (H), and number (n) of analyzed sections, are presented in Tab. 1. Overall, our quantification provides strong evidence that the preservation of tonotopic organization after adult-onset deafness correlates with the duration of hearing experience and thus with the age at deafness. Although they differed significantly from NHs rats in AVCN and CIC, OAD rats (∼12 months hearing + 1 month deaf) more or less maintained their tonotopy, whereas YAD rats (∼3 months hearing + 1 month deaf) interestingly lost their established organization already one month after deafness.

**Fig. 6:**
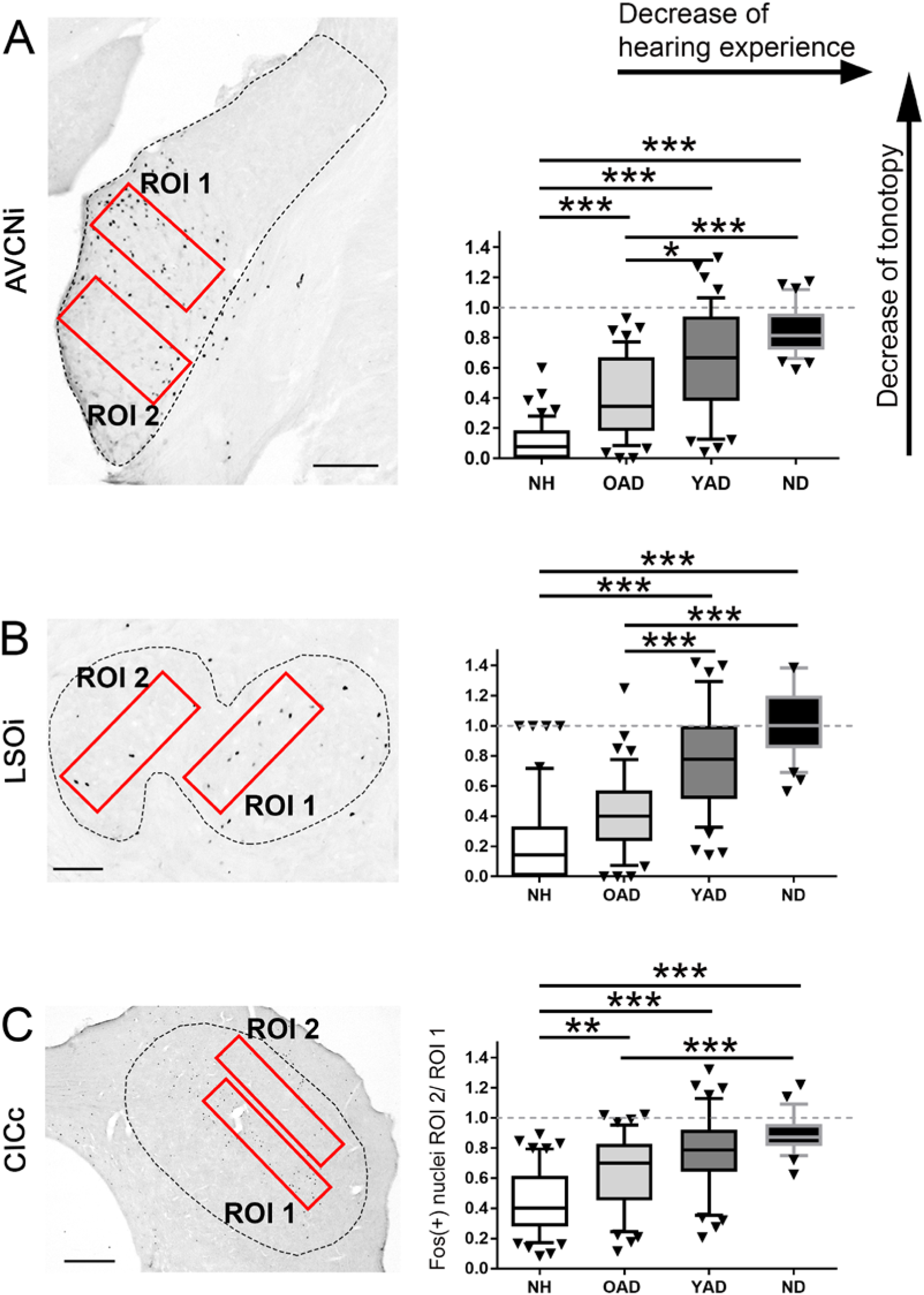
Quantification of the spatial distribution of the stimulation-induced Fos expression in the auditory brainstem of the four experimental groups, normal hearing (NH), old adult-deafened (OAD), young adult-deafened (YAD), and neonatally-deafened (ND) rats. A-C: Left figures show the two regions of interest (ROIs in red) in AVCNi (A), LSOi (B), and CICc (C) and the figures on the right show the results of the ratio calculation presented as box plots for each of the four cohorts. ROI 1 is placed in the mid-frequency range of a NH rat corresponding to the intracochlear stimulation position, ROI 2 covers the low frequency region of a NH rat. For statistics, the quotient of ROI 2/ROI 1 was calculated (right figures A-C). A small ratio corresponds to a good tonotopic order; a value of 1 means a Fos expression beyond the mid-frequency range of NH rats and thus a degradation of tonotopic order. Significant differences are shown by asterisks with * for p≤0.05, ** for p≤0.01, *** for p≤0.001. Box plots show distribution-based metrics of the 10th and 90th percentile and the median. AVCNi: anteroventral cochlear nucleus ipsilateral to electrical stimulation; CICc: central inferior colliculus contralateral to electrical stimulation; LSOi: lateral superior olive ipsilateral to electrical stimulation. Scale bars in A and B = 200 µm, and in C = 500 µm.

## 4. Discussion

A better understanding of the plastic changes that occur in the mature auditory system following acute deafness is an essential step towards improved rehabilitation of hearing-impaired patients and could contribute to improved auditory perception after the fitting with hearing prostheses. In order to contribute to this, the present study investigated the impact of deafness on the tonotopic organization of the mature auditory brainstem as a function of age at onset of deafness. One of the most important findings was that the central auditory system of younger and older adult rats responds differently to sudden and severe deafness. Using histological staining of the neuronal activity and plasticity marker Fos, we demonstrated for the first time that just one month of bilateral deafness results in a loss of tonotopic order in the auditory brainstem of younger rats, whereas the auditory system of older rats showed no comparable changes in the stimulation-induced Fos expression pattern. This suggests that the tonotopic order, as observed in normal hearing peers, is more or less preserved in older rats despite acute deafness.

### 4.1 Hearing function and induction of deafness in adulthood

Regardless of age, all of our female rats showed normal hearing thresholds prior to pharmacological deafening in adulthood. This observation coincides with a study on female Wistar rats by Möhrle et al. (2016), in which significant hearing loss could only be detected at the age of 19 months or older. Our 12-month-old animals showed hearing thresholds comparable with the thresholds of middle-aged animals (6-10 months) observed by Möhrle et al. (2016). This is different from the observations made for male Wistar rats, where an increase in hearing threshold is already observed at an age of 12 months and even earlier in noise exposed animals (Alvarado et al., 2014; Alvarado et al., 2019).

In addition to (Liu et al., 2011), we were able to demonstrate ototoxic efficacy of the combined compounds EA and KM not only for young adult rats (∼ 3 months old) but also for older rats (∼12 months old), as evidenced by the loss of the Preyer’s reflex (see Jero et al., 2001) and a strong increase in hearing thresholds by ∼92-93 dB SL, indicating the efficacy of this systemic deafness method regardless of the age of the animals. Despite pharmacologically induced deafness, which according to Liu et al. (2011) is mainly caused by irreversible degeneration of hair cells, both groups of EA and KM treated animals showed well differentiated EABRs following cochlear implantation. Under identical stimulation conditions and parameters, these EABRs were comparable to the EABRs of our normal hearing rats (see also Rosskothen-Kuhl et al., 2018; Jakob et al., 2019), indicating good and sufficient preservation of spiral ganglion neurons after one month of deafness.

### 4.2 Loss of tonotopy in younger but not in older auditory systems after acute deafness in adulthood

The expression of the protein Fos (also known as c-Fos) reflects the stimulation-induced activity of neurons (Morgan and Curran, 1986; Clarkson et al., 2010). Thirty to forty-five minutes of sustained electrical stimulation of the cochlea is sufficient to induce neuronal Fos expression along the auditory pathway of both NH as well as ND animals (Rosskothen-Kuhl and Illing, 2010, 2012). While frequency-specific stimulation of the cochlea of NH rats results in a local Fos expression pattern corresponding to the intracochlear stimulation position and thus reveals a tonotopic organization of the auditory pathway (Rosskothen et al., 2008; Jakob and Illing, 2009; Rosskothen-Kuhl and Illing, 2010; Jakob, 2011; Jakob et al., 2019), comparable stimulation of ND rats leads to a massively increased number of activated neurons, which are spread over almost the entire frequency range of the different auditory brainstem regions and thus indicates a loss of tonotopy (Rosskothen-Kuhl and Illing, 2012; Jakob et al., 2015; Rauch et al., 2016; Rosskothen-Kuhl et al., 2018; Jakob et al., 2019).

In this study, we could demonstrate that not only neonatal deafness but also acute deafness in young adulthood can result in a degraded tonotopic organization (Figs. 2-5). An auditory deprivation of one month, which corresponded to a quarter of the animal’s lifetime, was sufficient to degenerate the frequency mapping of the auditory pathway developed in the first three months of life as indicated by a spread of Fos(+) neurons beyond the “normal” frequency borders. By directly comparing the stimulation-induced Fos expression levels and patterns of all four cohorts, NH, OAD, YAD, and ND rats, we derived two main conclusions: First, the shorter the duration of deafness, the better the tonotopy of the auditory pathway, and second, with identical duration of deafness in adulthood, as in our YAD vs. OAD cohorts, age at onset of deafness and/or hearing experience appears to influence the preservation of tonotopy. In accordance with our results of the Fos quantification shown in Figure 6, a higher age at onset of deafness and thus a longer period of auditory experience, as in the case of our OAD cohort, seems to have a positive effect on the preservation of the tonotopic organization in the adult auditory system. In contrast to the older animals, the auditory system of younger animals seems to react quickly to the sudden absence of sensory input.

#### 4.2.1 Preservation of tonotopy in the older auditory system

Several studies have already shown that the plasticity level is higher in younger than in older brains (Berardi et al., 2004; Bavelier et al., 2010; Kwok et al., 2011; Sorg et al., 2016; Foscarin et al., 2017). An important role is attributed to the perineuronal nets (PNNs) in the central nervous system. PNNs are part of the extracellular matrix and play an important role in neuronal protection (Suttkus et al., 2012), the stabilization of synaptic contacts (Hockfield et al., 1990) as well as the inhibition of structural and functional plasticity (Frischknecht et al., 2009). In principle, the brain’s extracellular matrix mediates structural stability by enwrapping synaptic contacts fundamental, for example, for long-term memory storage (Happel et al., 2014). In the aging auditory system of rats, Mafi et al. (2020) have shown that the density of PNNs is massively increasing. Especially in the central and dorsal inferior colliculus a strong increase of PNNs on GABAergic and non-GABAergic cells could be identified. In addition, Foscarin et al. (2017) have demonstrated in rats that brain aging changes the sulfation of proteoglycans in PNNs, making the perineuronal nets more inhibitory and thus leading to a decrease in plasticity. A lower plasticity potential in the auditory system of our OAD cohort could be a reason why the tonotopic organization is still preserved after one month of deafness. Presumably, PNN-mediated inhibition of plasticity slows down or even prevents disinhibition of the auditory network in older rats, which can normally be triggered within hours after both juvenile and adult hearing loss (Browne et al., 2012; Llano et al., 2012; Resnik and Polley, 2017; Balaram et al., 2019). This hypothesis is in line with a study by Bavelier et al. (2010) who claim that “… a reduction in plasticity as development proceeds is likely to allow greater adaptability of the organism to variable conditions early in life, while ensuring an efficient neural architecture for known conditions by adulthood”. Whether the tonotopic organization in the OAD rats is still maintained even after a longer period of deafness or is lost with a delay cannot yet be answered and requires a follow-up study on OAD rats with a longer period of deafness. In contrast to the widespread and accepted hypothesis that aging can be associated with a reduced or even complete loss of plasticity (Kwok et al., 2011; Sorg et al., 2016; Foscarin et al., 2017; Persic et al., 2020), a study by Cisneros-Franco et al. (2018) has been shown that passive sound exposure can result in a plastic reorganization of the tonotopic map in the auditory cortex of older (age 22-24 months) but not younger adult rats (age 6-8 months). The increased but dysregulated plasticity was associated with a reduced inhibition of the aging auditory system. However, it should be noted that Cisneros-Franco et al. (2018) refer to cortical and not to subcortical structures and performed the experiments on hearing and not on deafened animals, thus limiting the applicability of these results to our study.

#### 4.2.2. Loss of tonotopy in the young adult auditory system

In contrast to the hypothesis that older auditory networks may have a lower plasticity potential, Heusinger et al. (2019) demonstrated a rapid degradation of the extracellular matrix after acute unilateral deafness in young adult rats (age 2-5 months). This was accompanied by an ingrowth of immature synapses into the AVCN of the deaf side. Furthermore, a modulation of the extracellular matrix in the CIC contralateral to deafness was observed already one day after deafness. This study thus provides important evidence that the young adult-deafened auditory system has a high plasticity potential and undergoes a remodeling of synaptic connections within only a few days after deafness due to short-term degradation of the extracellular matrix. According to these observations, it can be assumed that the auditory system of our YAD cohort was also still plastic and underwent remodeling as a result of acute hearing loss. A well-described consequence of adult deafness is changes in inhibitory networks. While studies on the auditory brainstem have identified inhibitory changes within days (Bledsoe et al., 1995; Abbott et al., 1999; Vale and Sanes, 2002; Vale et al., 2003), cortical networks undergo disinhibition within a few hours, as indicated by a reduced expression of GABAergic markers (Kotak et al., 2005; Sarro et al., 2008; Takesian et al., 2010; Browne et al., 2012; Llano et al., 2012; Takesian et al., 2012; Mowery et al., 2015; Resnik and Polley, 2017; Balaram et al., 2019). This could be one reason for the dramatic increase in neuronal activity and the accompanying degraded tonotopy in the auditory brainstem of our electrically stimulated YAD rats. In support of this hypothesis, a study by Gallinaro and Rotter (2018) on structural plasticity based on firing rate homeostasis in recurrent neuronal networks provide evidence that connectivity in sensory networks changes depending on stimulation and that a disturbance of the steady state could result in the degradation of connections, for example due to the loss of sensory input.

In agreement with our observations on young-adult rats, studies on other animal models and humans have shown that adult deafness can affect the tonotopic organization of the auditory pathway. Using fMRI on young to middle aged human patients, Wolak et al. (2017) demonstrate that postlingual sensorineural hearing loss affects the tonotopic organization of the auditory cortex. Significant differences in the cortical tonotopic map compared to normal hearing controls were also found in patients with bilateral high-frequency hearing loss (Koops et al., 2020). In line with this, the study by Robertson and Irvine (1989) demonstrates a plasticity of frequency organization in the auditory cortex of adult guinea pigs as early as 35 days after unilateral deafness. In addition, mild noise-induced hearing loss in adulthood leads to a broader frequency tuning in the primary auditory cortex of cats (Seki and Eggermont, 2002). According to Pienkowski and Eggermont (2011), deafness caused by damage to the auditory periphery induces plasticity in the adult auditory cortex, which is reflected in changes in the topographic map, among other things. In addition to the cortex, the auditory thalamus also shows plasticity of the tonotopic organization after unilateral deafness of adult cats (Kamke et al., 2003). In contrast to the auditory cortex and the medial geniculate body in the thalamus, no or only limited and non-permanent reorganization of the tonotopic map has been observed in regions of the auditory brainstem such as the dorsal cochlear nucleus or the CIC after deafness in adulthood (Rajan and Irvine, 1998; Irvine et al., 2003; Pienkowski and Eggermont, 2011). The previous observations for the auditory brainstem thus contrast with the results of our study in young adult-deafened rats, which for the first time demonstrated a deterioration of the tonotopy from the ventral cochlear nucleus via the LSO to the CIC of the auditory brainstem (even) after four weeks of auditory deprivation. Although the tonotopy of the auditory system has long been considered resistant to changes, e.g. hearing loss, after completion of a critical developmental phase, the above studies as well as our own results show that plasticity is preserved in the young adult brain and occurs at both cortical and subcortical levels.

### 4.3 Clinical relevance

As shown by Hiel et al. (2016), age is not a limiting factor for the CI fitting of patients. Even after a long period of deafness of 10-30 years a CI can still be of great benefit (Moon et al., 2014; Hiel et al., 2016; Medina et al., 2017). One possible explanation could be the increased stability of the hearing-experienced, mature auditory network in old age. Despite a reduced plasticity, a better preservation of network organization after deafness could help older patients to analyze sensory input via CI in a meaningful way. Provided that the data from the animal experiments can be transferred to human patients, our study provides evidence that elderly patients might be very suitable for reactivation of their auditory system by CIs due to the higher stability of their mature sensory networks even after a longer period of deafness. This hypothesis is supported, for example, by the better preservation of tonotopic organization after acute deafness in the old auditory system of mammals. This is in contrast to the auditory networks of younger patients, which are more plastic and therefore adapt more quickly to changes in sensory input. To date, the performance and benefit of older patients supplied with CI is still discussed controversially. While some studies show a better outcome for the younger adult CI patients (Chatelin et al., 2004; Vermeire et al., 2005; Lundin et al., 2013), others could not identify any differences (Labadie et al., 2000; Haensel et al., 2005; Mosnier et al., 2020; Bourn et al., 2022) or even demonstrate better performances for the elderly (Leung et al., 2005). For example, Leung et al. (2005) show that the age at implantation is only a low predictive value for postoperative performance. Elderly even performed better when the duration of deafness exceeded 25 years compared to their younger counterparts. Based on our data, we suppose that patients who become deaf at an advanced age might have a larger time window to benefit from CI supply in contrast to young adult patients who should be treated as soon as possible after deafening in order to counteract the degeneration of trained network organizations at an early stage. However, it is very likely that the latter, with appropriate training and rehabilitation, can compensate for a potential loss of network organization in the long term due to their higher level of plasticity.

### 4.4 Conclusion

The central auditory system of young and old adult rats adapts differentially to acute deafness in adulthood. Our study, in which the auditory system was (re)activated by electrical stimulation of the cochlea, demonstrated that the auditory pathway of young adult rats after acute deafness exhibits a loss of tonotopy comparable to that of neonatally-deafened rats. In contrast, after acute deafness in older animals, the auditory system retains its tonotopic order, similar to that of normal hearing animals. We conclude that neuronal plasticity predominates in the central auditory system of young adult rats. This is evidenced by the observation that acute deafness results in a degradation of frequency mapping along the auditory pathway after only a short period of deafness. In contrast, neuronal stability seems to prevail in the older auditory system. This could explain the preservation of frequency mapping in the auditory system even after a period of severe deafness and could be an optimal prerequisite for enabling a good hearing outcome after CI supply in older deaf patients.

## Data Access

The datasets used and analyzed during the current study are available from the corresponding author on reasonable request. All data generated or analyzed during this study are included in this published article.

## Acknowledgements

We thank H. Hildebrandt for technical assistance, proofreading, and helpful discussions and R.-B. Illing for continuous support and helpful discussion. Stimulation electrodes, programming software, and hardware components were kindly provided by Cochlear Germany GmbH & Co. KG, Hannover, Germany. We acknowledge support by the Open Access Publication Fund of the University of Freiburg.

## Authors’ contribution

NRK: Conceptualization, Resources, Data acquisition, Formal analysis, Supervision, Funding acquisition, Validation, Investigation, Visualization, Methodology, Writing - original draft, Project administration. TFJ: Data acquisition, Formal analysis, Investigation, Visualization, Writing-original draft. SG: Data acquisition, Formal analysis, Validation, Investigation, Methodology.

## Ethics approval and consent to participate

Animal experimentation: All procedures involving experimental animals reported here were approved by the Regierungspräsidium Freiburg (permission number G-10/83). All surgery was performed under ketamine (80 mg/kg) and xylazine (12 mg/kg) anesthesia, and every effort was made to minimize suffering.

